# Targeted-Neuroinflammation Mitigation Using Inflammasome-Inhibiting Nanoligomers is Therapeutic in Experimental Autoimmune Encephalomyelitis (EAE) Mouse Model

**DOI:** 10.1101/2024.01.11.575262

**Authors:** Sadhana Sharma, Sydney Risen, Vincenzo Gilberto, Sean Boland, Anushree Chatterjee, Julie A. Moreno, Prashant Nagpal

## Abstract

Multiple Sclerosis (MS) is a debilitating autoimmune disease that impacts millions of patients worldwide, disproportionately impacting women (4:1), and often presenting at highly productive stages of life. This disease affects the spinal cord and brain, and is characterized by severe neuroinflammation, demyelination, and subsequent neuronal damage, resulting in symptoms like loss of mobility. While untargeted and pan-immunosuppressive therapies have proven to be disease-modifying and manage (or prolong the time between) symptoms in many patients, a significant fraction are unable to achieve remission. Recent work has suggested more targeted neuroinflammation mitigation through selective inflammasome-inhibition can offer relief to patients, while preserving key components of immune function. Here we show a screening of potential therapeutic targets using inflammasome-inhibiting Nanoligomers (NF-κB1, TNFR1, TNF-α, IL-6), that meet or far-exceed commercially available small-molecule counterparts like Ruxolitinib, MCC950, and Deucravacitinib. Using the human brain organoid model, top Nanoligomer combinations (NF-κB1+TNFR1: NI111, and NF-κB1+NLRP3: NI112) were shown to significantly reduce neuroinflammation, without any observable negative impact on organoid function. Further testing of these top Nanoligomer combinations in an aggressive Experimental Autoimmune Encephalomyelitis (EAE) mouse model for MS using intraperitoneal (IP) injections showed that NF-κB1and NLRP3 targeting Nanoligomer combination NI112 rescues mice without observable loss of mobility or disability, minimal inflammation in brain and spinal cord histology, and minimal to no immune cell infiltration of the spinal cord and no demyelination, similar to or at par with mice that received no EAE injections (negative control). Mice receiving NI111 (NF-κB1+TNFR1) also showed reduced neuroinflammation compared to saline (sham) treated EAE mice and at par/similar to other inflammasome-inhibiting small molecule treatments, although it was significantly higher than NI112 leading to subsequent worsening clinical outcomes. Furthermore, treatment with an oral formulation of NI112 at lower doses showed a significant reduction in EAE severity, albeit with higher variance owing to administration and formulation/fill-and-finish variability. Overall, these results point to the potential of further development and testing these inflammasome targeting Nanoliogmers as an effective neuroinflammation treatment for multiple neurodegenerative diseases, and potentially benefit several patients suffering from such debilitating autoimmune diseases like MS.

## INTRODUCTION

### Multiple Sclerosis (MS) is an autoimmune neurodegenerative disease with inadequate treatment options

Approximately one million Americans and more than 2.8 million worldwide suffer from MS,^1,2^ an autoimmune demyelinating disease of the central nervous system that leads to irreversible disability. Given the high rate of diagnosis between the ages 20-50 (considered a formative and productive time for careers and families^3^), a disproportionate number of women that have MS (3.13 times more likely than men), and higher incidence rate in specific geographical locations, MS has an outsized impact on individuals affected, their families, and our society.^2–4^ MS occurs due to non-specific innate immune activation that creates a neurotoxic cocktail of inflamed microglia and astrocytes and attacks myelin, the insulation that covers and protects the nerves, leading to loss of motor function and disability. While there are no current cures for MS, disease-modifying therapies (DMT) focus on immune suppression, including corticosteroids, interferon beta suppression, and more recently CD20-directed antibodies to prevent B-cell activation. However, all DMTs have significant side effects with no neuroprotection and often also leave the individuals susceptible to other infections or immune diseases. More targeted therapeutics are being recently sought after, to provide better outcomes and longer disability milestones.^5^ While recent stem-cell therapies are directed towards neuroprotection and repair,^6^ there is a need for additional therapeutic options with a more targeted mechanism of action so as to reduce non-specific immune activation, prevent demyelination, and promote neuronal repair.

### Activated Inflammasome is a Novel Target for MS Therapy

Various parts of the immune system are involved in MS disease development and progression including inflammatory signaling pathways and activation of immune cells. Critically, CD4+ T cells, specifically Th1 and Th17 cells,^7–9^ play a crucial role in the inflammatory response in MS through recognition of myelin antigens, leading to the release of pro-inflammatory cytokines such as interferon-gamma (IFN-γ)^10–13^ and interleukin-17 (IL-17).^14–18^ Infiltrating immune cells release pro-inflammatory cytokines, including tumor necrosis factor-alpha (TNF-α), IL-1β, and IL-6, that contribute to inflammation and tissue damage within the brain and spinal cord. Another key part of the immune response is the activation of the complement system, part of the innate immune response, that leads to even further inflammation and tissue damage in MS patients.^19^ Axonal damage is another critical aspect of the neurodegenerative process in MS. The common inflammatory signaling events that are known to amplify or accelerate this disease pathogenesis seen in MS are mediated by NF-κB1 (nuclear factor kappa-light-chain-enhancer of activated B cells)^20,21^ and the activation of the NLRP3 (NOD-like receptor family, pyrin domain containing 3) inflammasome. Abnormal activation of the inflammasome, specifically NLRP3, has been actively investigated and associated with a wide range of inflammatory, autoimmune, cardiometabolic, and neurodegenerative diseases.^22–26^ Inflammasome inhibition has been intensely investigated as a novel target, as well as a common cure for a range of neurodegenerative and autoimmune diseases,^23,27–31^ specifically MS.^32,33^

### Nanoligomers Offer a Safe and Targeted Approach for Neuroinflammation Mitigation

Nanoligomer therapeutic platform generates gold nanoparticle-bound peptide nucleic acid (PNA) molecules to regulate any desired gene in a safe, effective, and targeted manner.^34–39^ PNAs demonstrate strong hybridization and specificity to their targets compared to naturally occurring RNA or DNA,^40^ and they exhibit no known enzymatic cleavage, leading to increased stability in human blood serum and mammalian cellular extracts.^41^ These advantages make PNAs a promising choice for expression-modifying therapeutics. Nanoligomers designed to inhibit the translation of target mRNAs simply bind the mRNA which prevents access to ribosomes. Given the low K_D_ (∼5 nM^37,39,40^), high binding specificity, minimal off-targeting,^37,39^ high safety profile, lack of any immunogenic response or accumulation in first pass organs, absence of any observable histological damage to organs even for long (>15-20 weeks) treatments,^37–39^ and facile delivery to the brain to counter neuroinflammation,^34,38^ make Nanoligomers and specifically NI112 cocktail a safe and targeted approach to mitigate neuropathological microglial and astrocytic activation.

## RESULTS AND DISCUSSION

### In-vitro Screening and Testing in Donor-Derived Primary Human Astrocytes and Human Brain Organoids

Since the inflammasome consists of a large number of potential targets up- and downstream, a target screen was done using reactive and activated astrocytes (or type A1) astrocytes.^42^ Briefly, activated microglia that are responsible for neurodegenerative diseases, have been shown to increase reactive A1 astrocytes via the release of pro-inflammatory cytokines. Using published protocols, the primary donor human astrocytes were activated using a stimulation cocktail of IL-1α, TNF α, and C1q (complement protein) to mimic the neurotoxic inflammation and induce an A1 phenotype (**Fig. 1A**).^42^ Following the stimulation of A1 astrocytes, as a high-throughput screen, top Nanoligomers targeting different inflammasome genes were added, and expression of key proinflammatory cytokine IL-1α was used to rank the impact of selected Nanoligomers on the downregulation of different inflammasome targets (**Fig. 1B**). The gene expression data showing gene down-regulation, and corresponding protein expression has been presented in prior publications.^34,35^ We collected multiplexed ELISA for 65 cytokines and chemokines (**Fig. S1**), and used Principal Component Analysis to analyze the key/combined outcome from these 65 biomarkers.^34,35^ Using Principal Component 1 which had accounted for more than 72% variance (PC1, 72.24%), we observed NF-κB1 and IL-6 as the two key protein targets that significantly reduce neuroinflammation (**Fig. 1C**). Further assessment of Nanoligomer molecules with corresponding small-molecule counterparts like Ruxolitinib, MCC950, and Deucravacitinib, revealed that the Nanoligomers meet or far-exceed commercially available therapeutic molecules targeting inflammation (**Fig. 1D**). Since IL-6 is involved in a number of biological functions in immunity, tissue regeneration, and metabolism,^43–47^ due to potential for side-effects in further screening, our algorithm^34,35^ identified NF-κB1 as the top target. We will conduct additional safety-toxicity assessment using animal models,^37–39^ before prioritizing and further testing of IL-6 inhibiting Nanoligomer. Therefore, after screening several targets to identify key nodes involved in the downregulation of neurotoxic chronic inflammasome activation, NF-κB1 was identified as the key single target that could restore innate immune response to the non-activated state. This is highly significant because so far NF-κB has been considered an “undruggable” target,^48,49^ and these results further highlight the advantages of the Nanoligomer targeting approach as opposed to traditional small-molecules and antibody-based therapeutic discovery. Further, other potential targets that could be synergistic with this key transcription factor, NF-κB1, were screened: 1) Combination of NF-κB1 and TNF receptor (TNFR1) downregulator (SB_NI_111 or renamed as NI111) ^34^; and 2) Combination of NF-κB1 and NLRP3 inflammasome downregulator (SB_NI_112 or renamed as NI112).^38^ These were termed as Nanoligomer combinations NI111 and NI112 respectively, and subsequently screened in the human brain organoid model (*in vitro*) and EAE model for MS (*in vivo*), to assess their pre-clinical efficacy for MS treatment, and identify molecular mechanisms that can be neuro- and myelin-protective.

**Fig. 1.**
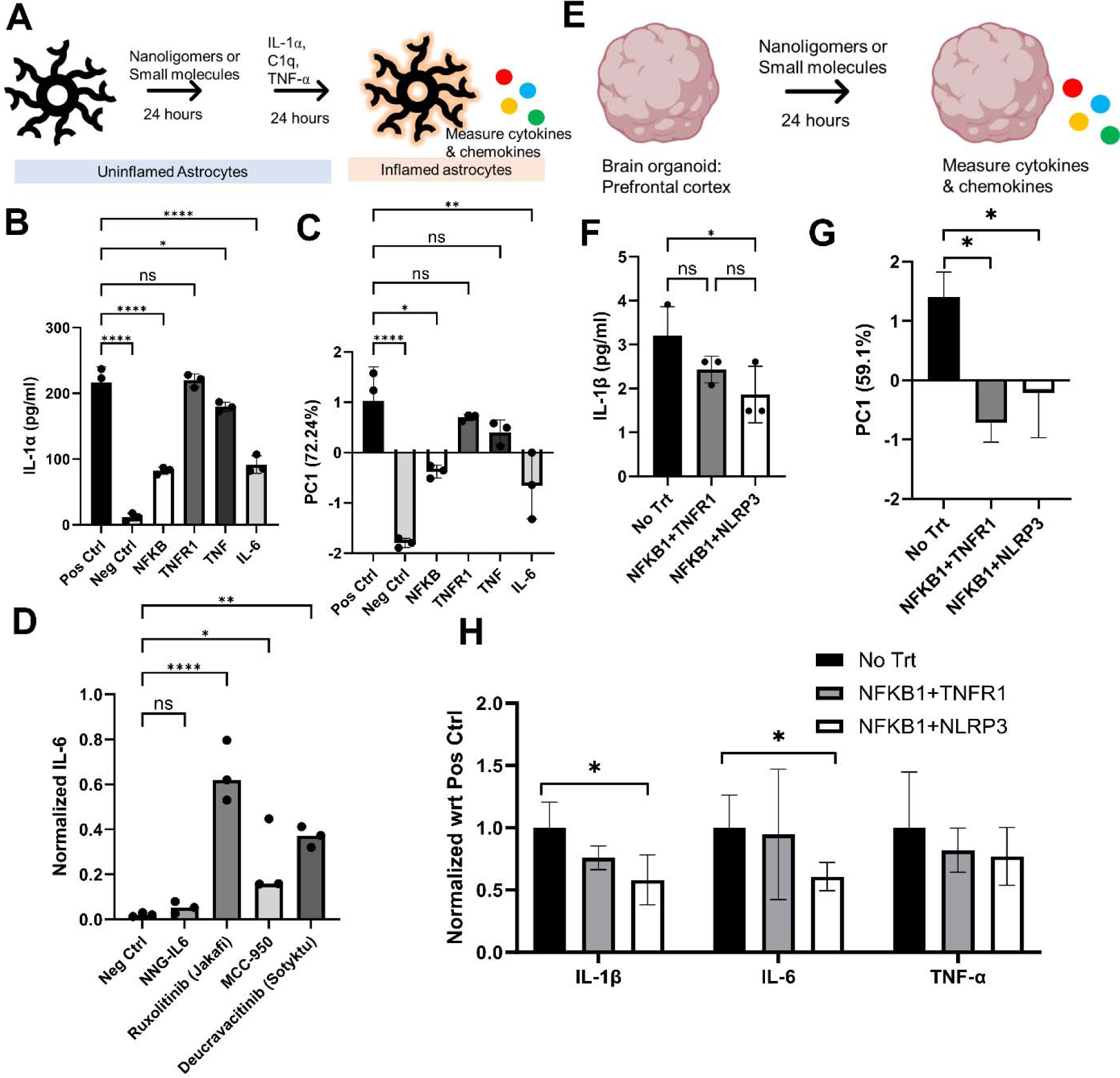
In vitro screening and ranking of single Nanoligomer targets, combinations, and comparison with small molecules, using donor-derived astrocytes and brain organoids. **A.** Schematic showing *in vitro* treatment and screening using activated astrocytes. **B.** IL-1α protein expression by untreated and treated astrocytes, used for screening and ranking of selected inflammasome-targeting Nanoligomers. **C.** Multiplexed 65-cytokine and chemokine expression (**Fig. S2**), was analyzed using Principal Component Analysis. PC1 (72.4% variability) was used for ranking the Nanoligomers. **D.** IL-6 cytokine expression, normalized by positive control (Pos Ctrl is 1), compared between IL-6 targeting Nanoligomer and respective IL-6/inflammasome targeting small molecule counterparts. **E.** Schematic showing *in vitro* screening using human organoids. **F.** Testing and ranking of NF-κB1 combination Nanoligomers using IL-1β. **G.** Ranking of Nanoligomers using combined 65-plex cytokine and chemokine expression using primary principal component PC1 (59.1%). **H.** Selected cytokines showing targeted downregulation of inflammasome by Nanoligomers (all 65-analysis shown in **Fig. S2**). \**P* < 0.05, \*\**P* < 0.01, and \*\*\**P* < 0.001, \*\*\*\**P* < 0.0001, Mean ± SEM, significance based on one-way ANOVA. *n* = 3 for each group.

### Screening top Therapeutic Combinations in brain organoid model

Following the identification of two synergistic combinations upstream and downstream with NF-κB1, NI111 and NI112 were assessed in human brain organoids. Briefly, human donor-derived iPSCs differentiated into different brain cell types (glutamatergic, dopaminergic, and gamma-aminobutyric acid (GABA)-ergic neurons), and astrocytes were grown together in brain organoids (to assess the impact on the prefrontal cortex (PFC) and ventral tegmental area (VTA) regions) and organoids^50^ (**Fig. 1E**). We assessed synaptic function using calcium imaging, followed by treatment with NI111 and NI112, to assess key inflammatory markers such as IL-1β (**Fig. 1F**) and 65-cytokine and chemokine biomarkers (**Fig. S2**). As before,^34,35^ we used the key Principal component PC1 (59.1%) and IL-1 as the screening and ranking method, to identify NI112 as the top therapeutic combination (**Fig. 1G**), followed by NI111. As seen here, both lead candidates, especially NI_112, showed significant downregulation in most biomarkers (**Fig. S2**) like IL-1β and IL-6 (p<0.05), except for TNF-α (**Fig. 1H**). These results showed potential for reversing neurodegenerative damage by reversing inflammasome activation and specific downregulation of key targets.

### Testing NI111 and NI112 in EAE mice model

We first validated the mechanism of action for translational inhibition of targeted genes by the Nanoligomer combination using immunoblotting (**Fig. 2A**). We homogenized brains of NI112 Nanoligomer-treated and sham-treated mice to confirm the reduction of activated NF-κB (through phosphorylated or phospho-p65 expression) and NLRP3 protein (p-value <0.05, see Methods for detailed protocol). EAE is the most commonly used experimental model for MS. EAE is a complex condition, where autoimmune demyelination is modeled through the administration of myelin basic protein peptide, in which the interaction between a variety of immune- and neuro-pathological mechanisms leads to an approximation of the key pathological features of MS: inflammation, demyelination, axonal loss, and gliosis.^51^ The counter-regulatory mechanisms of resolution of inflammation and remyelination also occur in EAE, which, therefore can also serve as a model for these processes. Briefly, Experimental Autoimmune Encephalomyelitis (EAE) induction was utilized to model multiple sclerosis. MOG_35-55_ antigen was delivered subcutaneously to induce a T-cell response, then an injection of the immune stimulator pertussis toxin, followed by another injection of MOG_35-55_ antigen (Hooke Labs). To address if Nanoligomers were neuroprotective in this mouse model of EAE we had the following four groups of female 10-week-old C57Bl6/J mice, 1.) untreated mice, 2.) EAE treatment + vehicle, 3.) EAE treatment + NI111 and 4.) EAE treatment + NI112. We verified the mechanism of action (MoA) and high safety profile of these Nanoliogmers in previously published studies.^34,52^ Treatments occurred three times a week at 150 mg/Kg of Nanoligomer or vehicle through intraperitoneal (IP) injection. Using the assessment of key motor function (motion of tail or limpness, partial or complete paralysis of hind limbs, partial or complete paralysis of front limbs, death, etc.) clinical scores are assigned to animals to evaluate MS progression. Clinical signs were monitored and recorded daily. Once a mouse had hind limb and partial front limb paralysis for two days mice would be scored a clinical score of 4 and culled and tissue taken. Mouse clinical scores were monitored for 14 days until scores of the vehicle mice reached four, at which time the study commenced, and tissue and blood samples were taken from all mice irrespective of their score (**Fig. 2B**). Typically, after 9-days, the MS progression is advanced with obvious inflammation and gliosis in spinal cord including demyelination, and progressively advancing paralysis and death. 13 days after EAE induction, >2 animals reached clinical scores requiring euthanasia (>4 for more than 2 days, or score of 5) as per institutional IACUC protocols (Colorado State University), and the experiment was terminated.

**Fig. 2.**
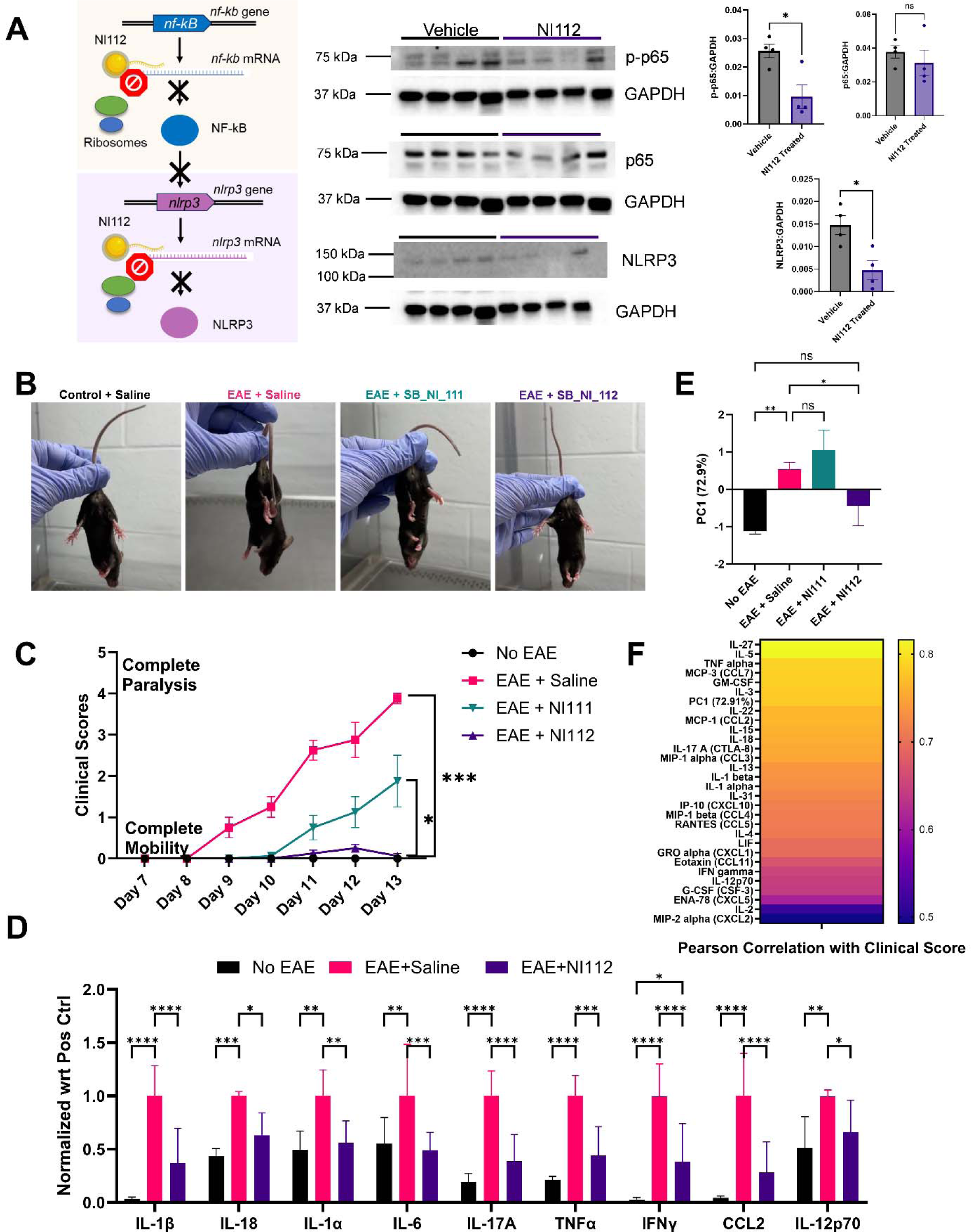
Nanoligomer NI112 significantly reduces neuroinflammation, reduces EAE severity and prevents disability. **A.** (Left) Schematic showing the mechanism of action, (Right) Immunoblotting images and quantification showing a clear reduction in activated NF-κB (or phosphorylated-p65, p-p65) and NLRP3 in NI112 treated mice brains. Expression levels of p65 unit of NF-κB is unchanged. All proteins are normalized with GAPDH. **B.** Representative images of clinical signs of EAE including limp tail and hind leg inhibition. Images taken at 11 days post-immunization (11 dpi). **C.** Mean clinical scores from 7 dpi to 13 dpi. Clinical scoring key is provided in the methods. Higher clinical scores show increased disability, whereas low scores (or zero) show protection from disease. **D**. Selected key cytokine and chemokines measured in homogenized spinal cord tissue of No EAE, EAE + Saline, and EAE+NI112 group mice, using multiplexed ELISA. **E.** Principal Component 1 (72.9% variance) was used for assessing different treatment groups. Negative numbers indicate lower inflammatory cytokine and chemokine expression (lower neuroinflammation). NI112 treatment mimics no EAE (negative control). **F**. Heatmap showing calculated Pearson Correlation between clinical scores and PC1 and inflammatory cytokine and chemokines measured for each mouse as a replicate, across all treatment groups. \**P* < 0.05, \*\**P* < 0.01, and \*\*\**P* < 0.001, \*\*\*\**P* < 0.0001, Mean ± SEM, significance based on one-way ANOVA. *n* = 4-8 for each group (n=4 no EAE, EAE + Saline, n=8 for NI111 and NI112 treatment). Method of administration: intraperitoneal (IP) injection.

As seen here (**Fig. 2B, C, Supplementary EAE Videos**), while NI111 delayed the disease progression and was statistically significant in disease modification (as DMT) like other inflammasome and NLRP3 therapies such as Dapanustrile/OLTT1177^32^ and MCC950^33^, demyelination still occurred leading to paralysis (no evidence of disease activity or NEDA is not achieved). NI112 on the other hand was able to successfully achieve clinical scores identical to no EAE control (**Fig. 2C, Supplementary EAE Videos**). We would like to note that such an observation has not been reported by other small molecule inflammasome inhibitors^32,33^ and DMTs, which nominally show a delay in onset and timing of disease progression, but not complete halting of disease advancement, as reported here. Further, biochemical and histological examination revealed no evidence of demyelination (**Fig. 3**). It is important to note that 2 mice started to show some evidence of limpness in the tail (clinical scores 0.5 on the scale of 5 on day 11), but the limpness was gone by day 13, indicating some potential for remyelination might have occurred. Histological examination through Luxol fast blue (LFB) staining showed no myelin loss in these or other mice treated with NI112.

**Fig. 3.**
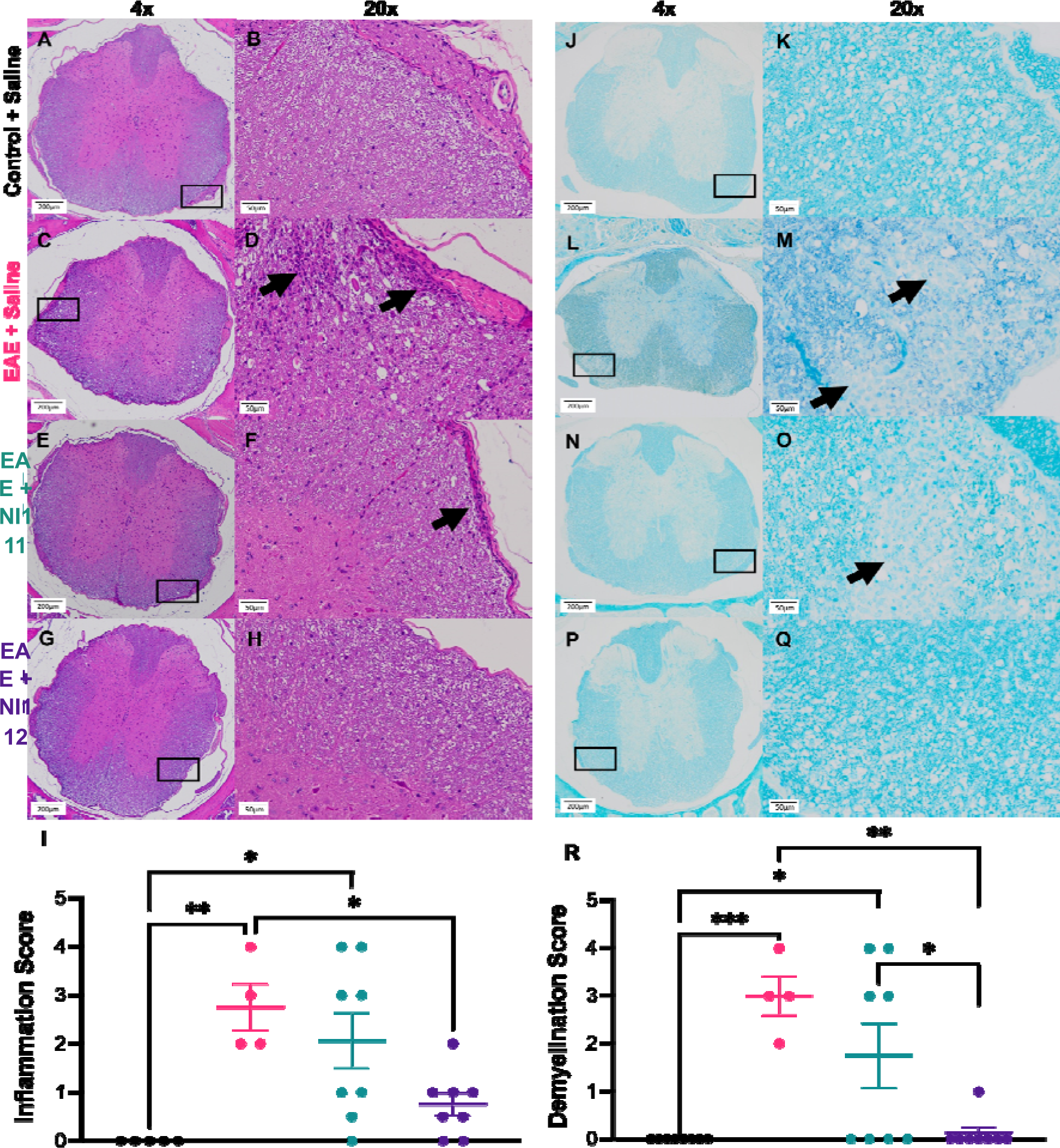
NI112 treatment significantly inhibits myelin loss and immune cell infiltration in EAE. **(A-H).** Representative images of hematoxylin-eosin (H&E) stained spinal cord at 4x (left) and 20x (right) of Control + Saline, EAE + Saline, EAE + NI111, and EAE + NI112 groups. Arrows identify areas of immune cell infiltration. **I.** Evaluation of inflammation score from H&E staining. **J-Q.** Representative images of Luxol fast blue (LFB) stained spinal cord at 4x (left) and 20x (right) of Control + Saline, EAE + Saline, EAE + NI111, and EAE + NI112 groups. Arrows identify areas of demyelination. **R.** Evaluation of demyelination score from LFB staining. \**P* < 0.05, \*\**P* < 0.01, and \*\*\**P* < 0.001, Mean ± SEM, significance based on one-way ANOVA. *n* = 4-8 for each group (n=4 no EAE, EAE + Saline, n=8 for NI111 and NI112 treatment). Method of administration: intraperitoneal (IP) injection.

### Biochemical assessment of the spinal cord tissue for NI112 treated mice in EAE model

Beyond clinical scores, the spinal cord and brain tissue of EAE-treated mice were assessed for inflammation and neurodegeneration biomarkers for MS. As seen in **Figs. 2D, 2E, S3**, in both the spinal cord, NI112 significantly decreased the inflammatory biomarkers. We assessed a significant increase of these levels in the spinal cord, and to a lesser extent, in the brain of EAE vehicle-treated mice. However, not only did the Nanoligomer lead treatment prevent paralysis, but the spinal cord (and brain tissues) also did not exhibit the upstream and downstream cytokines like IL-1β, IL-18 (directly linked to NLRP3), and IL-6, but also exhibited a decrease in other B-cell and T-cell activation chemokines like IL-17 (**Fig. 2D, S3**). Further, a recent assessment of specific biomarkers linked to primary progressive MS (PPMS, IL-1β, IL-6, TNF-α, CCL3, CXCL1, CXCL2, CXCL10, NLRP3)^53^ which has even fewer treatment options. NI112 shows a significant decrease of all these reported PPMS biomarkers and promise for advancing this Nanoligomer cocktail toward underserved MS patients (**Fig. 2A, 2D, S3).** All results point towards inhibition of microglial activation, to prevent these from attaching to the myelin sheath. To further strengthen the power of biochemical assay, we assessed the correlation of biochemical markers/immunological assay with functional/clinical scores for respective mice, across different groups. As can be seen using the Pearson Correlation Coefficient (**Fig. 2F**), we saw an extremely strong correlation (>0.5) for almost all cytokine/chemokine, and principal component 1, further validating the strength of conclusions drawn here and in vivo assessment. Further, histological analysis was conducted using H&E, IBA1, and Luxol (**Figs. 3,4**), to assess inflammation scores and potential demyelination, and confirmed that Nanoligomer NI112 treated mice showed no disease evidence activity.

**Fig. 4.**
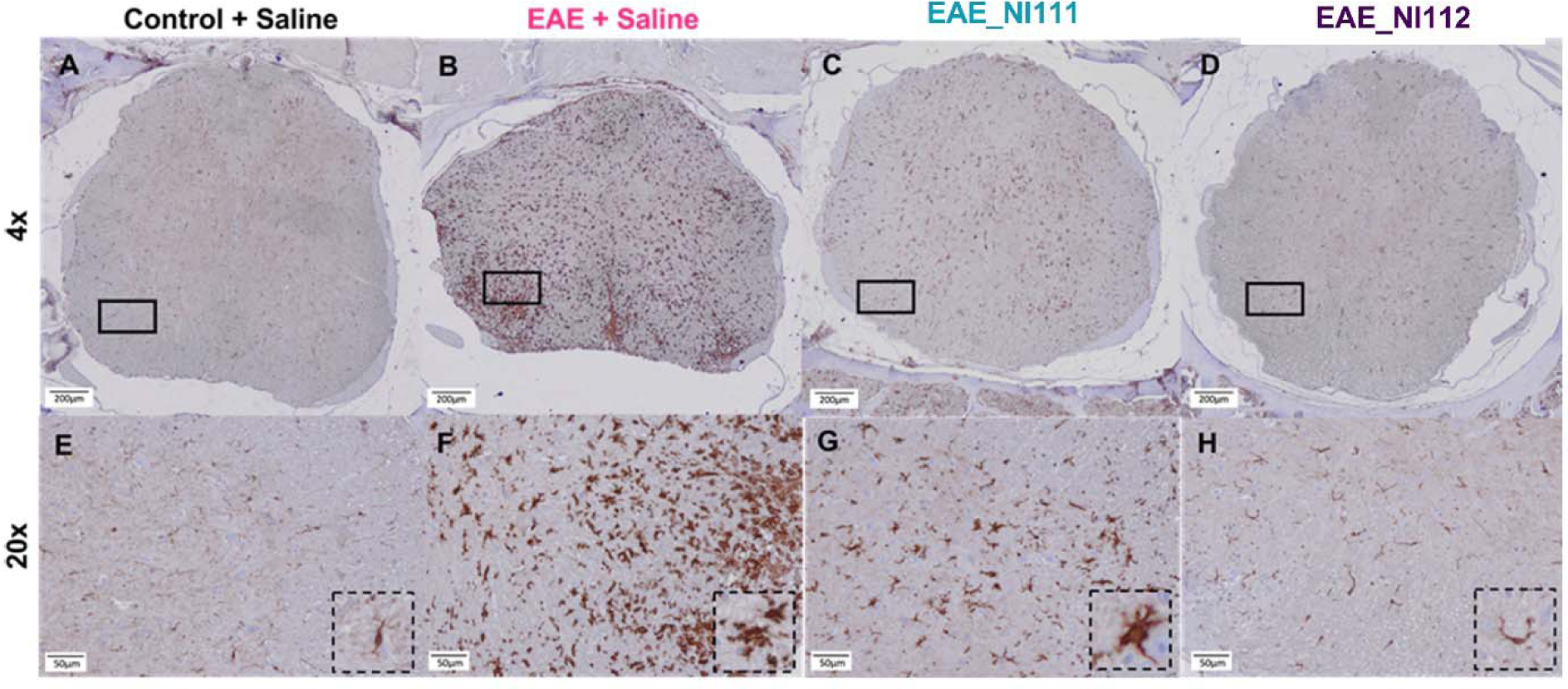
NI112 protects against microglial inflammation during EAE. **(A-H)** Representative images of Iba1 stained spinal cord at 4x (top) and 20x (bottom) of Control + Saline, EAE + Saline, EAE + NI111, and EAE + NI_112 groups. Solid rectangles represent 20x region. Dashed squares display the morphology of single microglia.

### Histology for Inflammation and Demyelination assessment in EAE model

Spinal cords were extracted from each of these mice and fixed in 10% normal buffered formalin for 48 hours followed by decalcification, paraffin wax processing, and embedding. Tissue was sliced at 5µm and stained with hematoxylin and eosin and imaged to detect filtration of immune cells into the white matter of the spinal cord. These spinal cords were then scored from 1-4 (with 4 being the most infiltration of immune cells), with a significant increase in immune cell filtration in mice exposed to EAE + vehicle and EAE + NI111 compared to control mice with no EAE induction. The EAE + NI112 were significantly decreased from the EAE + vehicle and not significantly different from the control-treated mice (representative **Figs. 3A-H**, inflammation scores in **I).** Next, we analyzed the myelination of spinal cords with Luxol Fast Blue Stain (representative **Figs. 3J-Q**), scoring the spinal cord images from 0-4 for the degree of demyelination, with 4 being the most demyelination and 0 having no demyelination (**Fig. 3R**). In the control mice (i.e. no EAE) and NI112 there was no loss of myelin and a score of 0 was given while mice treated with NI111 had no significant change when compared to the EAE + vehicle cohort (**Fig. 3R**). The reason for the lack of demyelination (and potential remyelination) can be seen from the lack of microglial activation using NI112 treatment, as seen using IBA-1 staining in spinal cords (**Fig. 4**).

### Oral Dosing using NI112 in EAE model

Besides injection-based delivery, we assessed an oral formulation of NI112 in the EAE model. Using Size M capsules (Torpac) with Eudragit (Evonik) enteric coating, we filled dried NI112 powder (maximum API packing 60-75 mg/kg) and administered the dose orally using an oral gavage applicator (Torpac). We observed that besides variability in capsule filling and coating (fill and finish), the administration introduced additional challenges with some mice biting or breaking the capsules during administration. We saw a bimodal distribution in clinical scores for dosed mice where more than half of the treated group of mice showed no signs of disease (like IP administration), whereas a minority group showed delayed progression/disease delay but clear onset and progression of typical EAE signs (**Fig. 5A**) like other inflammasome small-molecule therapeutics.^32,33^ Therefore, we assessed the drug amount using ICP-MS and showed clearly the mice that were properly dosed showed zero clinical scores and more than 3-fold higher drug concentration than mice that showed disease onset and lower drug concentration (**Fig. 5B**). Further assessment using tissue multiplexed ELISA showed similar neuroinflammation mitigation (**Fig. 5C, 5D, S4**) and strong Pearson Correlation (>0.5) between biochemical/immunological assessment and clinical scores (**Fig. 5E**). Histological assessment also validated lack of immune cell infiltration and lack of any myelin sheath loss in oral-dosed EAE mice (**Fig. 6**). These results show the flexibility in using different Nanoligomer formulations.

**Fig. 5.**
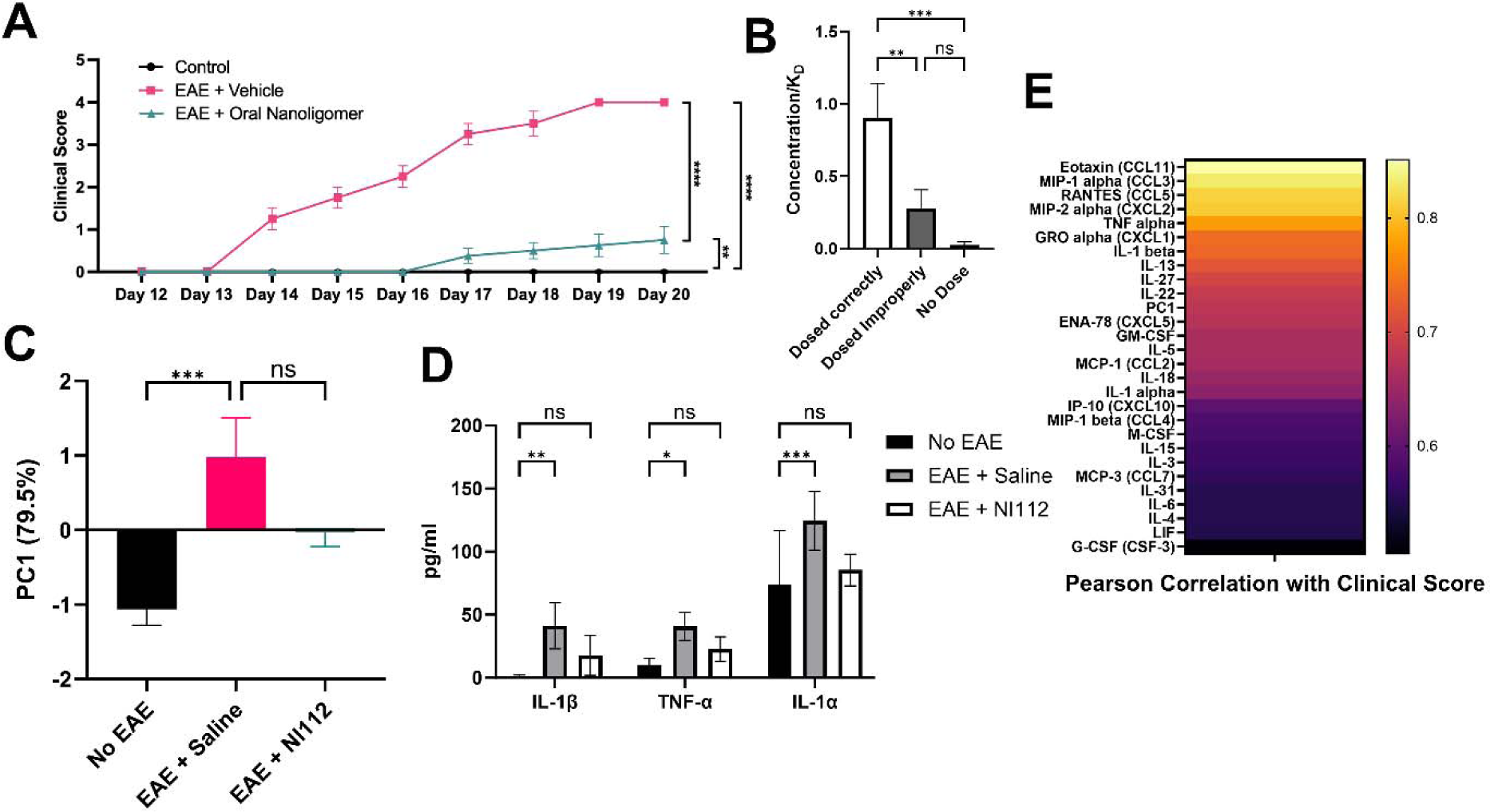
Oral dosing NI112 is therapeutic in the EAE mice model. **A.** Mean clinical scores showing improved neuroprotection and mobility in NI112 treated mice. **B**. Measured NI112 concentration (normalized by drug K_D_ 5nM^37,39,40^) showing significant variation in correctly and incorrectly dosed mice, providing evidence that only properly dosed mice show neuroprotection and low clinical scores. The observed variability was due to differences/lack of optimized protocol for oral administration and API capsule coating (fill and finish). **C**. Principal Component Analysis (PC1 79.5%) showing a large difference between NI112 treated and sham EAE mice. Entire 36-cytokine and chemokine analysis is shown in **Fig. S4**. The NI112-treated mice are closer to No EAE than untreated mice. **D**. Selected cytokine in homogenized spinal cords for mice in the three groups (No EAE or negative control group, EAE + Saline or positive control group, and EAE + NI112 or treatment group). **E**. Heat map of Pearson correlation coefficients between behavioral (clinical scores) and biochemical (36-cytokines and chemokine measured in spinal cord tissue) for all three groups of mice. \**P* < 0.05, \*\**P* < 0.01, and \*\*\**P* < 0.001, Mean ± SEM, significance based on one-way ANOVA. *n* = 4-8 for each group (n=4 no EAE, EAE + Saline, n=8 for NI112 treatment). Method of administration: enteric-coated oral capsule.

**Fig. 6.**
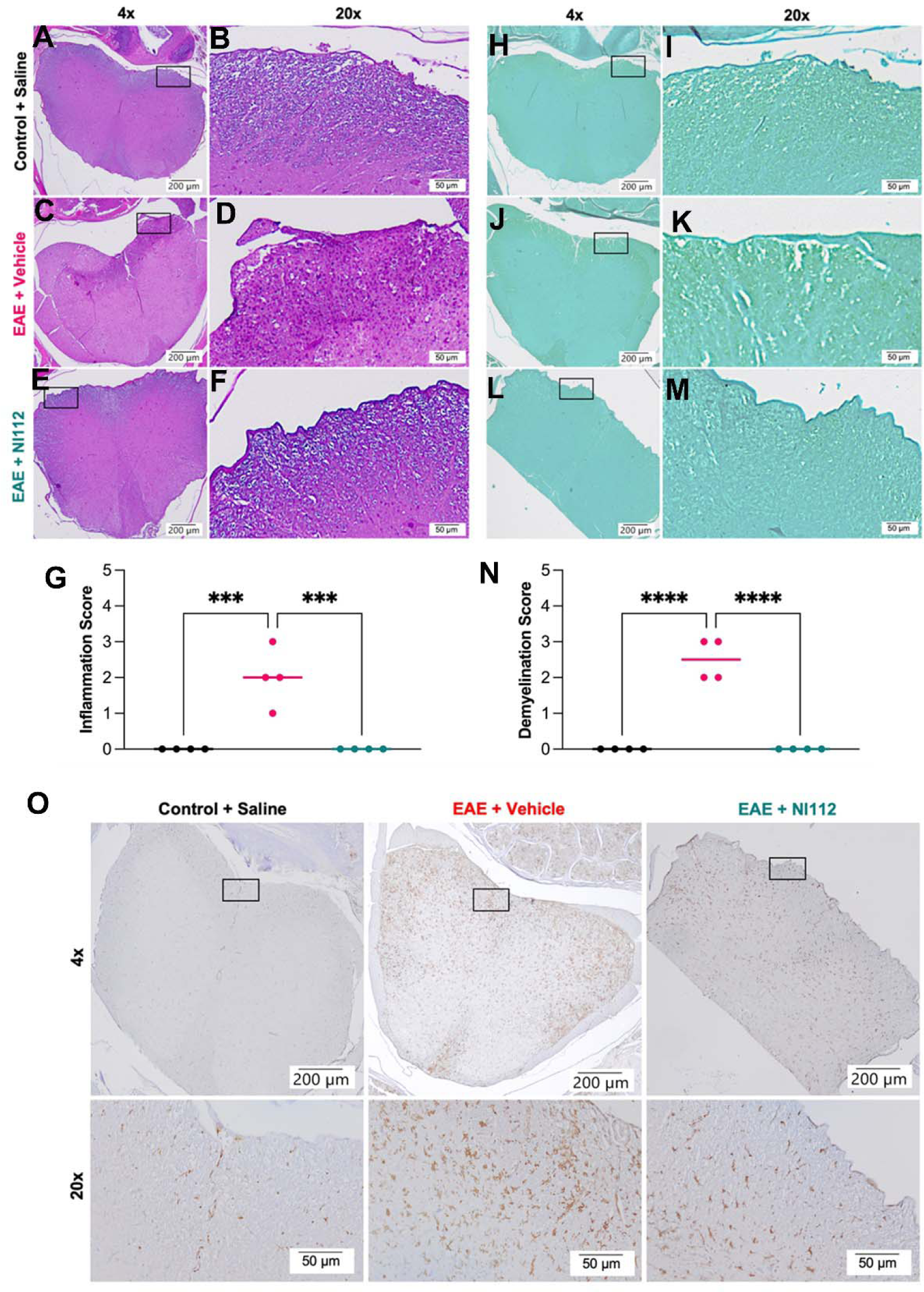
Inhibition of NF-κB1 and NLRP3 by oral treatment of NI112 significantly protects against microglial inflammation, and inhibits immune cell infiltration and myelin loss in EAE. **(A-F)** Representative images of hematoxylin-eosin (H&E) stained spinal cord at 4x (left) and 20x (right) of Control + Saline, EAE + Saline, and EAE + NI112 groups. Arrows identify areas of immune cell infiltration. **(G)** Evaluation of inflammation score from H&E staining. **(H-M)** Representative images of Luxol fast blue (LFB) stained spinal cord 4x (left) and 20x (right) of Control + Saline, EAE + Saline, EAE + NI111, and EAE + NI112 groups. Arrows identify areas of demyelination. **(N)** Evaluation of demyelination score from LFB staining. **(O)** Representative images of Iba1 stained spinal cord at 4x (top) and 20x (bottom) of Control + Saline, EAE + Saline, and EAE + NI112 groups (A-F). Solid rectangles represent 20x region. *p<0.05, **p<0.01, and ***p<0.005, Mean ± SEM, significance based on one-way ANOVA, n=2-8 for all groups.

## CONCLUSIONS

### Targeted Mitigation of Neuroinflammation induced pathology key for New MS Therapeutic Development

To replace pan immunosuppressive DMTs, more targeted therapies are being sought. Specifically, targeting inflammasome has been suggested as an important alternative, and our work showed screening of selected inflammasome targets using 65-cytokine and chemokines resulted in the identification of NF-κB1. Nevertheless, NF-κB1 has traditionally been deemed an “undruggable” target,^48,49^ emphasizing the advantages of the Nanoligomer targeting approach over conventional modalities such as small-molecules and antibody-based therapeutic discovery. Moreover, the specific combination of NF-κB1 with NLRP3 showed higher neuroinflammation mitigation compared to even other effective combinations (NF-κB1 and TNFR1^34^), adding another dimension to optimizing efficacy while preserving key immune functions.

### NF-**κ**B1 and NLRP3 Combination far Surpasses other reported small-molecule studies

As noted above, an effective NF-κB1 + TNFR1 combination NI111 showed appreciable neuroinflammation mitigation.^34^ As a result, NI111 delayed the disease progression in the EAE model and was statistically significant, as other DMTs and inflammasome and NLRP3 therapies such as Dapanustrile/OLTT1177^32^ and MCC950^33^, demyelination still occurred leading to paralysis. However, the NI112 combination (NF-κB1 + NLRP3) far surpassed these small molecules^32,33^ in neuroinflammation mitigation, as well as other EAE studies, showing therapeutic effect in the EAE model using both injection and oral formulation. While injection formulation showed consistent results in different *in vitro* and *in vivo* models, oral formulation showed larger variability, likely due to non-optimal administration and formulation fill-and-finish. Further human clinical translation is needed to advance and test these inflammasome targeting Nanoliogmers as an effective neuroinflammation treatment.

### Summary Conclusions

In summary, our study demonstrates that selectively inhibiting NF-κB1 and NLRP3 to mitigate targeted neuroinflammation can be therapeutic in the EAE mouse model. The Nanoligomer combination, NI112, effectively rescued mice, preserving mobility, preventing disability, minimizing inflammation in histology, and exhibiting minimal to no immune cell infiltration or demyelination. These outcomes were comparable to negative control mice without EAE injections and surpassed other small-molecule counterparts. We employed two formulations, utilizing IP injections and enteric-coated oral capsules. Both methods significantly reduced EAE severity, though oral dosing exhibited higher variance due to administration and formulation variability. These findings suggest the potential for further exploration and development of these targeted inflammasome Nanoligomers as a promising treatment for neuroinflammation, with the capacity to address various neurodegenerative diseases and potentially benefit individuals affected by debilitating autoimmune conditions such as MS.

## MATERIALS AND METHODS

### Nanoligomer design and synthesis

Nanoligomers were specifically designed and synthesized (Sachi Bio) following established methods detailed in published references.^34–39^ The Nanoligomers are composed of an antisense peptide nucleic acid (PNA)^34–36,54^ conjugated to a gold nanoparticle.^34–36^ The 17-base-long PNA sequence was selected to optimize solubility and specificity, targeting the start codon regions of *nfkb1* (Sequence: AGTGGTACCGTCTGCTA) and *nlrp3* (Sequence: CTTCTACTGCTCACAGG), within the Mouse genome (GCF_000001635.27), identified using our bioinformatics toolbox.^36^ Various PNA sequences were screened for their solubility, self-complementing sequences, and potential off-target interactions within the mouse genome (GCF_000001635.27). The PNA portion of the Nanoligomer was synthesized on a Vantage peptide synthesizer (AAPPTec, LLC) employing solid-phase Fmoc chemistry. Fmoc-PNA monomers, wherein A, C, and G monomers were shielded with Bhoc groups, were obtained from PolyOrg Inc. Post-synthesis, the peptides were attached to gold nanoparticles and subsequently purified using size-exclusion filtration. The conjugation process and the concentration of the refined solution were monitored by measuring absorbance at 260 nm (for PNA detection) and 400 nm (for nanoparticle quantification).

### Culturing and treating 3D organoids

We used already published protocols and methods to culture 3D organoids.^50,55^ Briefly, differentiated iCellNeurons (Fujifilm Cellular Dynamics) were thawed and mixed together in ultralow attachment (Sbio) plates and cultures for 3 weeks using complete BrainPhys media, until the organoids were matured. Typical PFC organoids used 90:10 Neuron: Astrocyte ratio with 70:30 distribution of glutamatergic: GABAergic neurons.^50^ Following the 3-week maturation period, we used 200 µM Nanoligomer stock and replaced 5% of BrainPhys media with stock. Untreated negative controls used stock media. Small molecule compounds Ruxolitinib, MCC950, and Deucravacitinib (Cayman Chemicals) were added at 10uM final concentration using respective stocks in PBS. Following 24-hour treatment, the media was exchanged and analyzed using multiplexed ELISA, using manufacturer recommendations, as detailed below.

### In vivo testing in EAE and control mice

10–13-week-old female C57Bl/6J mice were purchased from Jackson Labs. EAE induction was done using MOG_35-55_ antigen (Hooke Labs) delivered subcutaneously on Day 0 at the top of the spine and base of the spine. Two hours after they are given an intraperitoneal (IP) injection of the immune stimulator pertussis toxin PTX dose. 24 hours later (Day 1) another dose of PTX was administered IP. Nanoligomer cocktail (1x intraperitoneal injection; 150 mg/kg body weight) or vehicle was started on Day 2 (compared to treatments starting Day 0 for other inflammasome therapies^32,33^), then given every other day until the dissection date. Mice were housed throughout all experiments at ∼18-23°C on a reverse 12 light/12 dark cycle. Fresh water and *ad-libitum* food (Tekland 2918; 18% protein) was routinely provided to all cages. Animals were consistently health checked by the veterinary staff at Colorado State University (CSU). This protocol was approved by CSU IACUC and Laboratory Animal Resources staff.

### Tissue collection

After deep anesthetizing with isoflurane, ∼1 ml of blood was removed via cardiac puncture followed by cervical dislocation. The left hippocampus and a piece of left cortex tissue were removed, flash-frozen on dry ice, and stored at -80°C until further processing. The right hippocampus and right cortex were processed (paraffin-embedded) for immunostaining (details below). The spinal cords were also collected with the cervical and lumbar regions preserved in 10% normal buffered formalin (NBF) and the thoracic portion was flash-frozen on dry ice and stored at -80°C until further processing.

### Clinical scoring

Clinical scores were assigned daily using the following scheme.

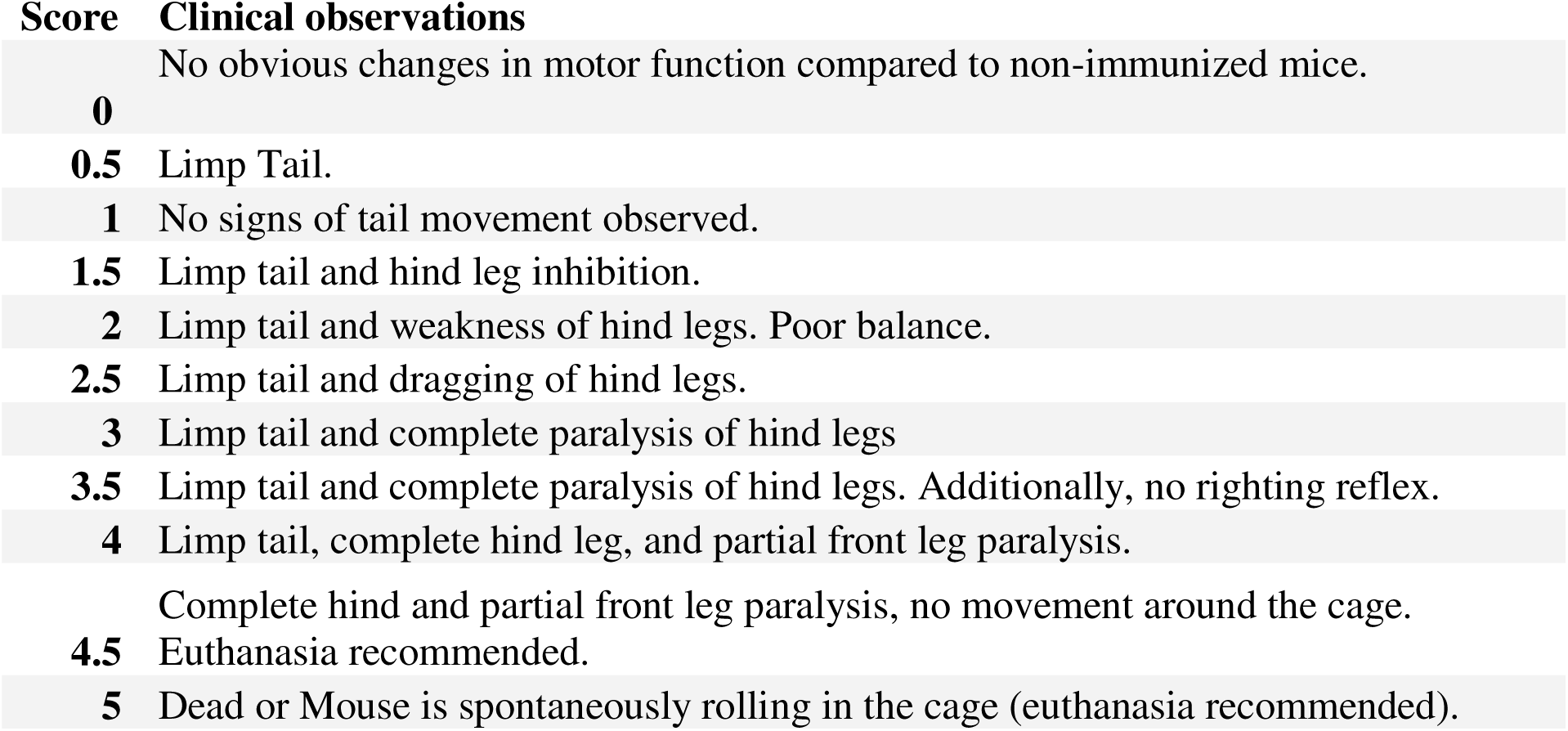

### Multiplexed ELISAs

Tissue homogenates from all mice underwent evaluation utilizing the ThermoFisher 36-plex Procartaplex cytokine/chemokine panel, following previously documented procedures.^34,35^ Briefly, 25uL of 10 mg/ml spinal cord or brain tissues were homogenized using ProcartaPlex™ Cell Lysis Buffer (ThermoFisher) using 5-mm stainless steel beads (Qiagen) at 25 Hz for 1.5-3min (Qiagen/Retsch Bead Beater). Following homogenization samples were centrifuged at 16,000 x g for 10 mins at 4 °C. After centrifugation homogenized samples were measured for protein concentration. Homogenate was then handled in accordance with the manufacturer’s provided protocol and analyzed using a Luminex MAGPIX xMAP instrument.). Standards for each cytokine/chemokine were employed at 1:4 dilutions (8-fold dilutions), alongside background and controls. Subsequently, concentrations of samples were determined from a standard curve using the Five Parameter Logistic (5PL) curve fit/quantification method.

### Immunohistochemistry and myelin staining

Paraffin-embedded spinal cords were sectioned at 4um before staining. Immune cell filtration into the spinal cords and morphological changes were examined pathologically by hematoxylin and eosin staining. Slides were deparaffinized and rehydrated before being treated with hematoxylin and bluing reagent. Tissue was counterstained with eosin (Epredia). To determine the degree of immune cell infiltration, images were scored from each cohort of mice using a score from 0 to 5, with 0 being no infiltration of immune cells to 5 being large pockets of infiltration of cells. Demyelination was scored using Luxol Fast Blue Stain Kit (Abcam) for myelin, with scores also ranging from 0 to 5, with zero being a perfectly myelinated spinal cord to 5 where most of the cord is demyelinated. For immunohistochemistry analyses of microglia/macrophage infiltration, paraffin-embedded tissue was sectioned at 4 µm, deparaffinized, rehydrated, sodium citrate treated, and blocked in Tris A/2% donkey serum (Jackson ImmunoResearch), then incubated overnight in the following IBA1 antibody (1:50 dilution; Abcam)Wash steps were performed using 2% bovine serum albumin and 2% Triton-X in 1LJM TBS, and an ABC HRP peroxidase detection kit (Vector Laboratories) and ImmPACT DAB Substrate Peroxidase (HRP) Kit (Vector Laboratories) were used for a chromogen. Slides were counterstained with hematoxylin (ThermoFisher), secured with a coverslip in mounting medium, and stored at room temperature until imaging. Whole tissue images were taken on an Olympus BX53 microscope with an Olympus DP70 camera using an Olympus UPlanSApo 20x objective (N.A. = 0.75). Representative images were taken using an Olympus BX53 microscope with an Olympus DP70 camera using an Olympus UPlanFL N 40x objective (N.A. = 0.75).

### Immunoblotting

Four saline/sham-treated and four NI112-treated C57Bl/6 (Jackson Laboratory) mice were intracranially inoculated with 30µl of 0.1% 22L or Rocky Mountain Laboratories (RML) strains of mouse-adapted prions. After harvesting brain tissue, mouse brain homogenates were created by utilizing lysis buffer (50mM Tris, 150mM NaCl, 2mM EDTA, 1mM MgCl2, 100mM NaF, 10% glycerol, 1% Triton X-100, 1% Na deoxycholate, 0.1% SDS and 125mM sucrose) supplemented with Phos-STOP and protease inhibitors (Roche). The protein concentration of the lysates was determined using a BCA Protein Assay kit (Thermo Scientific). Subsequently, the lysates underwent centrifugation at 40,000 x g for 1 hour at 4°C before being applied to a gel. The samples were electrophoresed on 4-20% acrylamide SDS page gels (BioRad) and then transferred to 0.45µm PVDF blotting paper (Millipore). Primary antibodies used include phosphorylated nuclear factor kappa B (p-NF-κB, 1:1000 dilution, ABclonal); nuclear factor kappa B (NF-κB; 1:1000, ABclonal), NLR Family Pyrin Domain Containing 3 (NLRP3; 1:1000, Cell Signaling). Membranes were incubated in primary antibody (TBS-T in 5% BSA) for 24 hours at 4°C, followed by HRP-conjugated secondary antibodies (TST-T in 5% milk; 1:5000; EMD Millipore). For loading control, GAPDH was run at a 1:5,000 dilution (Millipore), with HRP-conjugated anti-mouse secondary at 1:5,000 dilution (Southern Biotech). The visualization of the protein-antibody complex was achieved by employing the SuperSignal West Pico PLUS Chemiluminescent Substrate (Thermo Scientific) and imaged using the BioRad ChemiDoc MP. The quantification was done using ImageJ Software.

### Oral formulation

Size M (mice) oral capsules (Torpac) were completely filled with dried Nanoligomer NI112 mixed with dicalcium phosphate (LFA) in a 50:50 ratio. The weighed capsules (∼3mg/capsule) had 75mg/kg API dosing for 20 mg mice. The capsules were then coated with Eudragit (Evonik) enteric polymer, dried, and given to mice using an oral gavage applicator (Torpac). The resulting API concentration was measured using ICP-MS at the University of Colorado Anschutz facility, using previously published protocols.^37,39^

### Statistics

The figure legends specify the statistical tests conducted, the number of mice involved, and the reported p-values. To evaluate variances among groups, a one-way ANOVA with Tukey’s post-hoc testing was employed, except for a two-group comparison (Fig. 2A) where the unpaired t-test was used. Additionally, the Pearson correlation method was utilized to establish relationships between cytokine levels and cognitive function data. Microsoft Excel and GraphPad Prism 10.1.0 were employed for data analysis, and GraphPad Prism software was used for data presentation.

## ASSOCIATED CONTENT

### Supporting Information

Supporting figures S1-S4, with 65-plex Multiplexed human ELISA plots for *in vitro* screening, 36-plex cytokine and chemokine plots for spinal cord tissue ELISA in the EAE mouse model.

## Author Information

### Author Contributions

P.N. conceived the idea, designed the experiments, and synthesized the Nanoligomers. S.S. and V.G. conducted all the *in vitro* experiments and biochemical characterization. S.R., J.A.M., and S.B. conducted the EAE mouse studies. S.S., A.C., and P.N. conducted the ELISA measurements. P.N. and A.C. wrote the manuscript with input from all the authors. All authors read the manuscript and provided input.

### Notes

### Declaration of competing interests

S.S., V.G., A.C., and P.N. work at Sachi Bio, a for-profit company that developed the Nanoligomer technology. A.C. and P.N. serve as the founders of Sachi Bio, with P.N. having filed a patent on the technology. The remaining authors declare no competing interests.

## ACKNOWLEDGMENTS

Authors acknowledge financial support from NASA SBIR Awards 80NSSC22CA116 and 80NSSC23CA171.

## TOC GRAPHIC

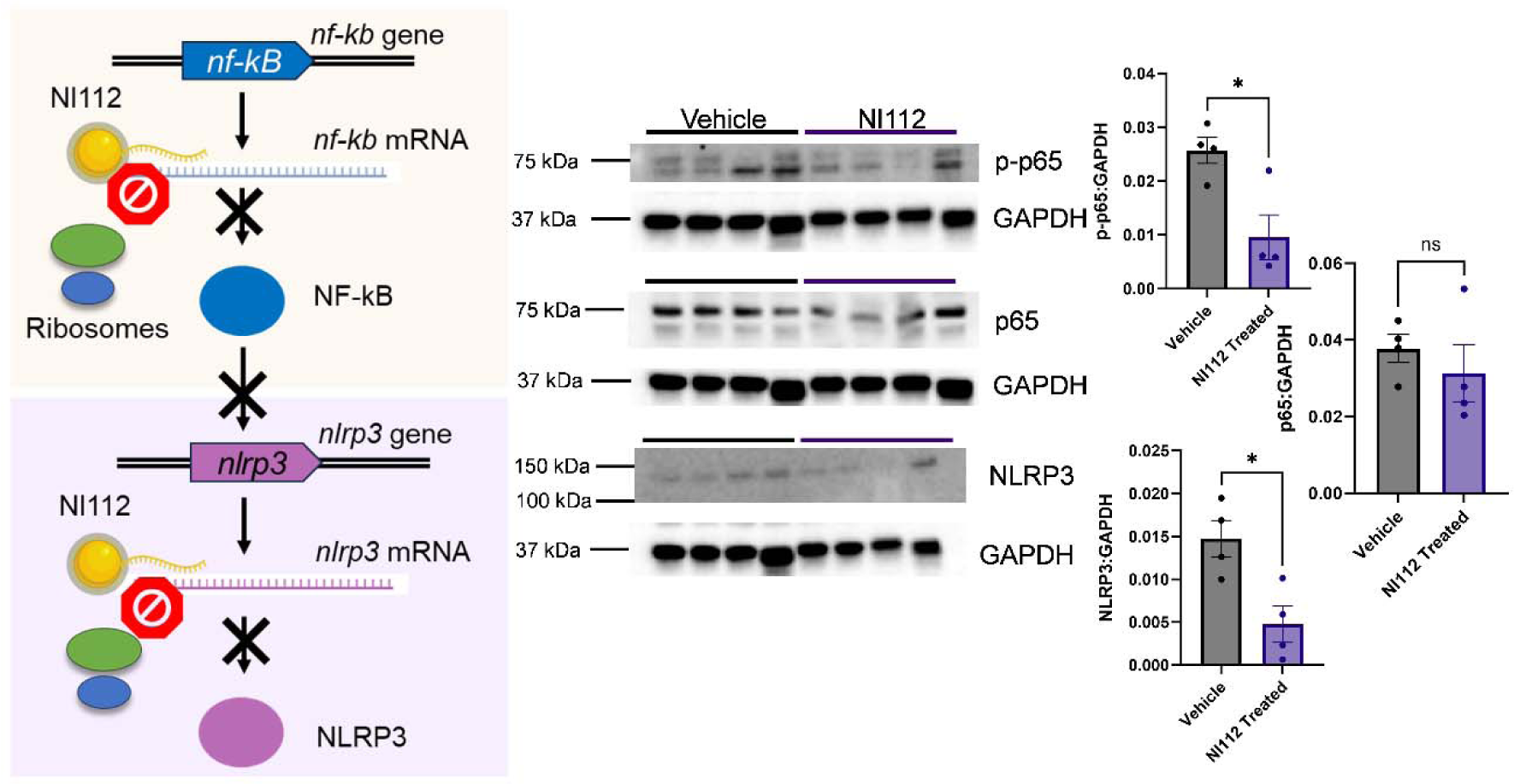

## Notes

### Summary of Updates

Figure 2 and 5 revised, text added to highlight biomarkers related to primary progressive MS

