## Supporting Information for "Targeted-Neuroinflammation Mitigation Using Inflammasome-Inhibiting Nanoligomers is Therapeutic in Experimental Autoimmune Encephalomyelitis (EAE) Mouse Model"

**Supplementary Figures:**

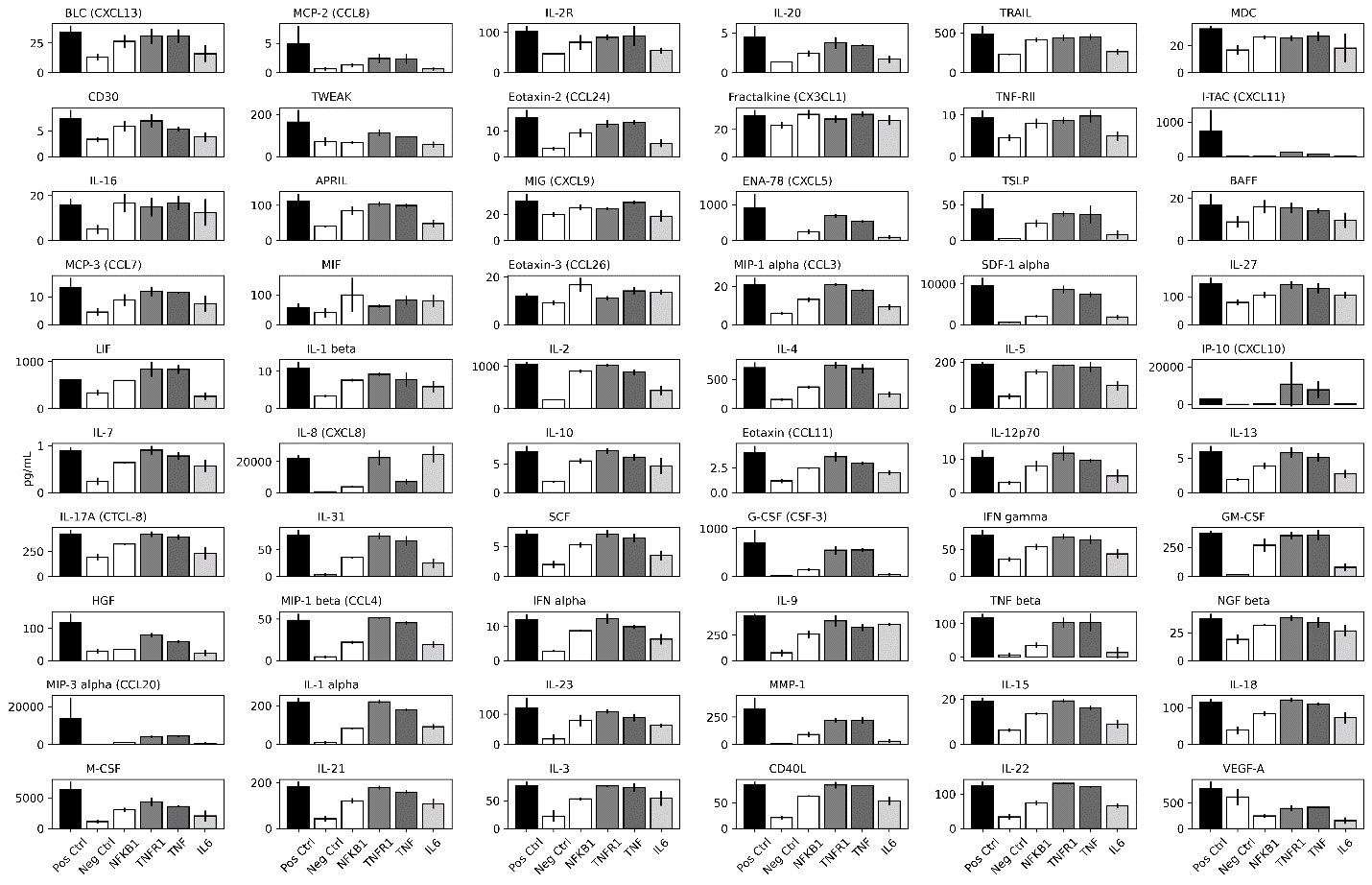

**Fig. S1. Screening inflammasome targets using 65-plex Multiplexed ELISA analysis of human cytokines and chemokines.** Donor-derived human astrocytes were stimulated using cytokine cocktail (IL-1α, TNF- α, C1q) and then treated with respective top Nanoligomer molecules for different inflammasome targets (NFκB1, TNFR1, TNF, IL-6) when compared to Positive and Negative Controls. The following cytokines were not plotted here: IL-6, FGF-2, TNF-α, MCP1 (CCL2), and GROα (CXCL1), due to saturation (upper limit of detection). Some of these (e.g., IL-6 shown in **Fig. 1D**) were subsequently measured using individual Simplex kits (ThermoFisher) for direct comparison of different treatments.

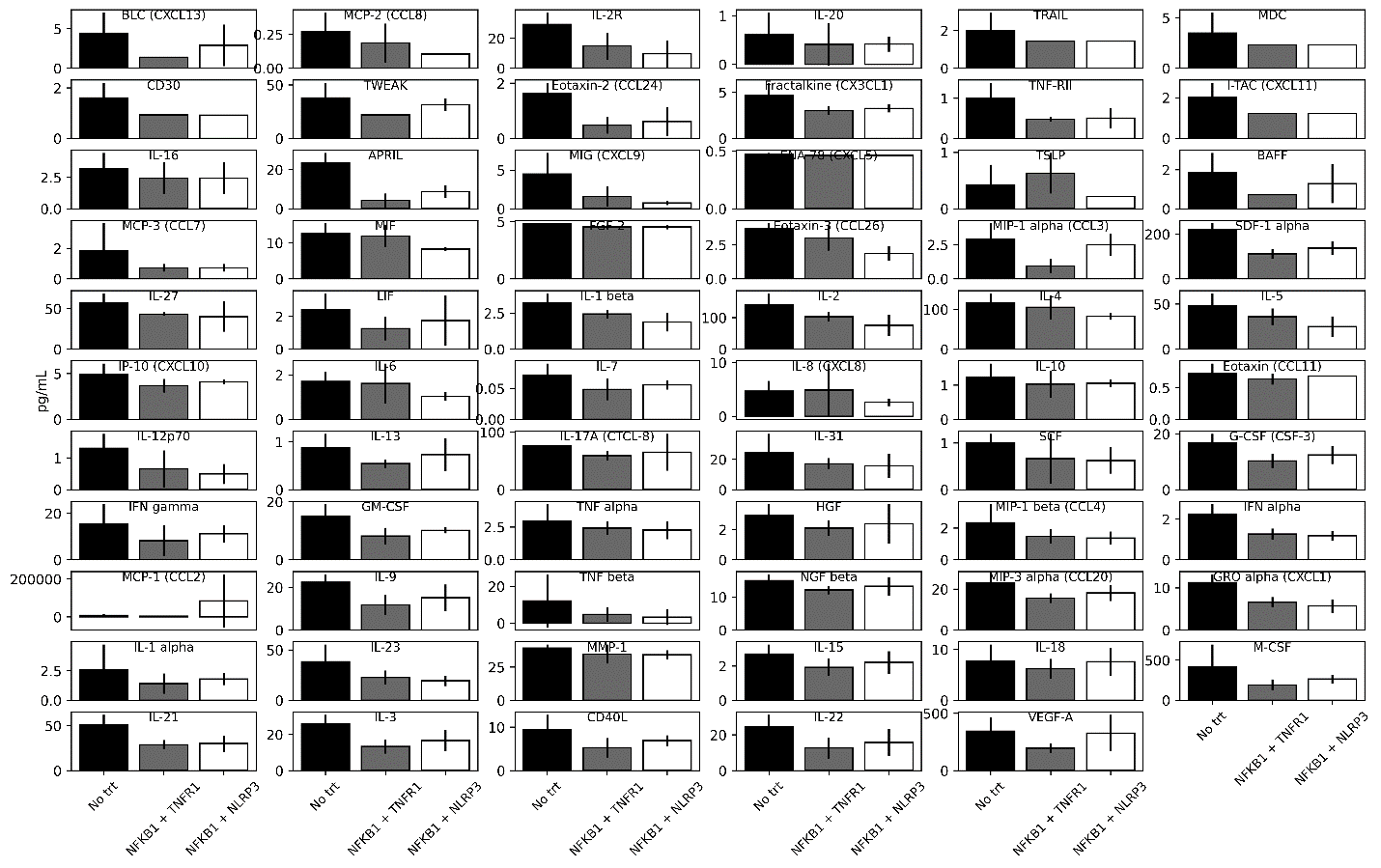

**Fig. S2. Testing top Nanoligomer target combinations (NFκB1+TNFR1 and NFκB1+NLRP3) in human brain organoids, using 65-plex Multiplexed ELISA analysis of human cytokines and chemokines.** Human brain organoids were treated with top Nanoligomer combination molecules (NFκB1+TNFR1, NFκB1+NLRP3). No treatment (Trt) shows negative control.

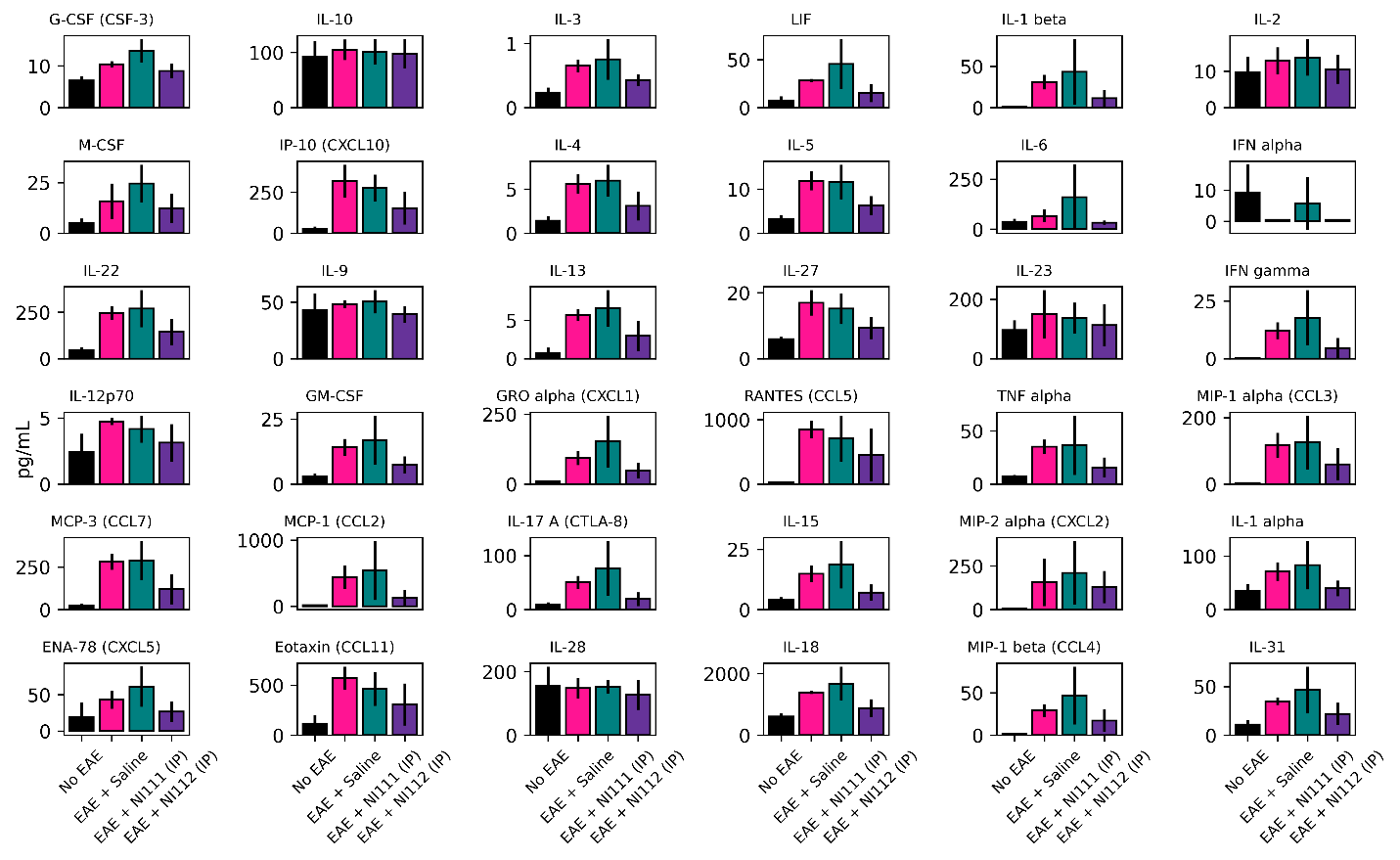

**Fig. S3. Comprehensive assessment of** **NI11 and NI112 (Mouse) Nanoligomer combinations using** **36-plex Multiplexed ELISA analysis of mouse cytokines and chemokines.** Mouse spinal cords were homogenized and assessed using Multiplexed ELISA. No EAE mice were non-immunized mice showing negative control, whereas EAE + Saline showed positive control group, for each cytokine and chemokine.

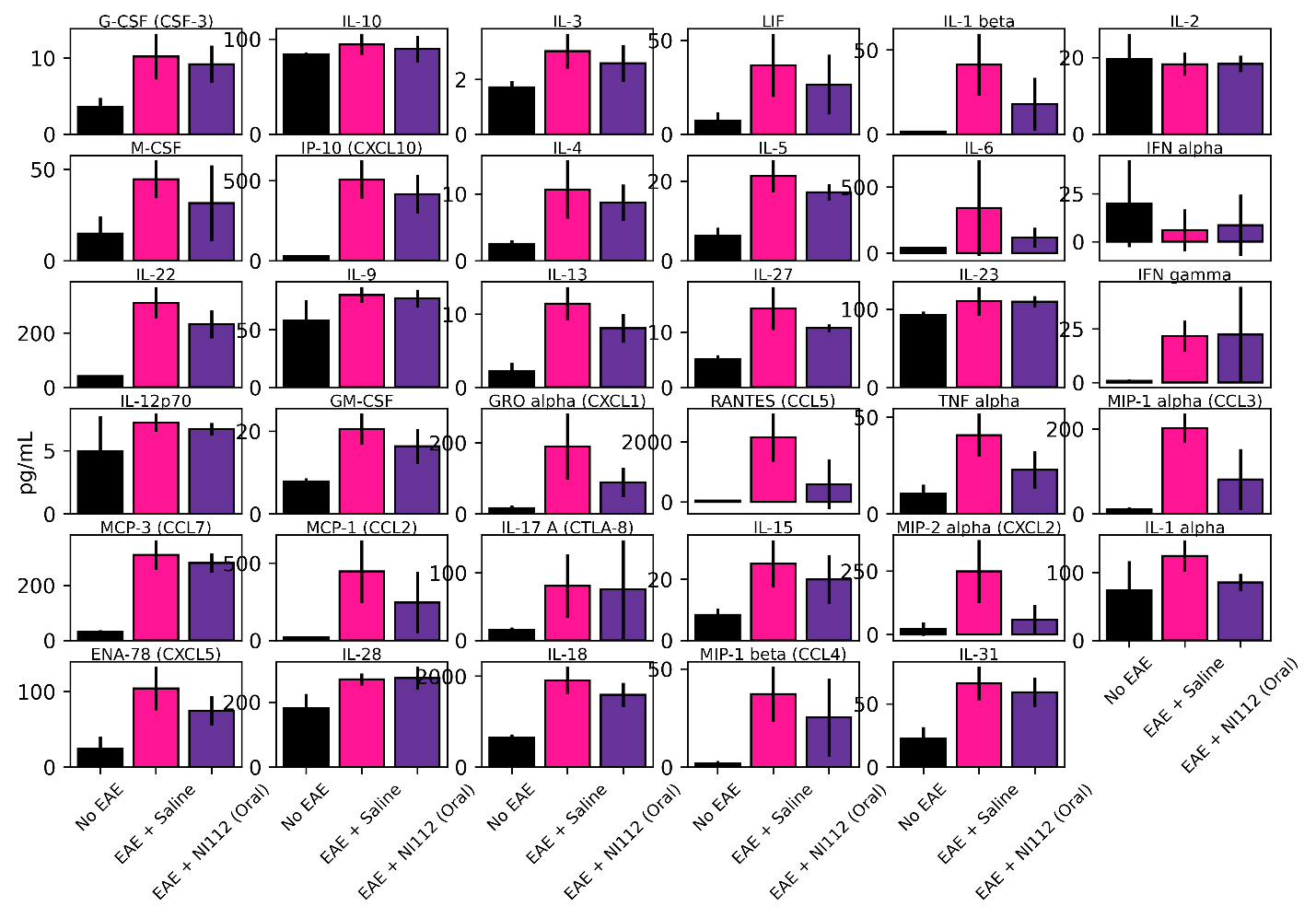

**Fig. S4. Comprehensive assessment of** **orally administered** **NI112 (Mouse) Nanoligomer combinations using** **36-plex Multiplexed ELISA analysis of mouse cytokines and chemokines.** Mouse spinal cords were homogenized and assessed using Multiplexed ELISA. No EAE mice were non-immunized mice showing negative control, whereas EAE + Saline showed positive control group, for each cytokine and chemokine. Eotaxin/CCL11 was not plotted here due to limit of detection.

**Supplementary EAE Videos**:

**Healthy Control**: Folder has 2 representative videos showing gait and tail movement for healthy (No EAE) mice. These are the negative control mice.

**EAE Vehicle**: Folder has 4 videos showing EAE + Vehicle 4 mice, with increasing paralysis and clinical scores. These are the positive Control mice.

**EAE SB_NI_111**: Folder has 8 videos showing EAE + SB_NI_111 8 mice, with lower inflammation than positive control, but increasing paralysis and clinical scores.

**EAE SB_NI_112**: Folder has 8 videos showing EAE + SB_NI_112 8 mice, with minimal inflammation or paralysis, low clinical scores, healthy mice at par with negative control (No EAE) mice.
